# Myelin Basic Protein dynamics from out-of-equilibrium functional state to degraded state in myelin

**DOI:** 10.1101/602441

**Authors:** M. Di Gioacchino, A. Bianconi, M. Burghammer, G. Ciasca, F. Bruni, G. Campi

**Affiliations:** Roma Tre University; Rome International Center for Materials Science Superstripes (RICMASS); ESRF; Physics Institute, Catholic University of Sacred Heart; Università di Roma Tre; CNR Institute of Crystallography

## Abstract

Living matter is a quasi-stationary out-of-equilibrium system; in this physical condition, structural fluctuations at nano- and meso-scales are needed to understand the physics behind its biological functionality. Myelin has a simple ultrastructure whose fluctuations show correlated disorder in its functional out-of-equilibrium state. However, there is no information on the relationship between this correlated disorder and the dynamics of the intrinsically disordered Myelin Basic Protein (MBP) which is expected to influence the membrane structure and overall functionality. In this work, we have investigated the role of this protein structural dynamics in the myelin ultrastructure fluctuations in and out-of-equilibrium conditions, by using synchrotron Scanning micro X Ray Diffraction and Small Angle X ray Scattering. We have induced the crossover from out-of-equilibrium functional state to in-equilibrium degeneration changing the pH far away from physiological condition. While the observed compression of the cytosolic layer thickness probes the unfolding of the P2 protein and of the cytoplasmic P0 domain (P0_*cyt*_), the intrinsic large MBP fluctuations preserve the cytosol structure also in the degraded state. Thus, the transition of myelin ultrastructure from correlated to uncorrelated disordered state, is significantly affected by the unfolding of the P2 and P0 proteins, which in this latter state do not act in synergistic manner with MBP to determine the membrane functionality.

**STATEMENT OF SIGNIFICANCE:** A better comprehension of myelin degenerative process and the role of protein dynamics in this biological membrane is a topic issue in today’s scientific community. The myelin ultrastructural fluctuations exhibit correlated disorder in its functional state, that becomes uncorrelated as it degenerates. In this work we elucidate the interplay of protein structural dynamics and myelin ultrastructure in the transition from its functional state to the degraded state. The results highlight that the intrinsically disordered Myelin Basic Protein (MBP) allows to preserve the myelin structure following both the small correlated fluctuations in physiological state and the large disordered fluctuations in degraded conditions, where the myelin functionality is close to being lost and the MBP remains the single active protein.

## INTRODUCTION

The protein kingdom consists of two main classes, the 3D structured proteins and the intrinsically disordered proteins (IDPs). This latter class of functional proteins, with highly dynamic fragments, is characterized by a lack or partial lack of ordered 3D structure in physiological condition. These do not follow the classical structure-function paradigm and then exist as heterogeneous ensembles of conformers (1). IDPs display a random-coil-like average conformation, studied as isolated polypeptide chain under physiological condition (2–4). However, the IDPs have a fundamental importance in many biological processes, such as fold on binding to their biological targets or constitute flexible linkers that have a role in the supramolecular configurations at nano- and meso-scales (5). In this context we chose to study the myelin basic protein (MBP) an IDP (6–9) that is involved in the self-assembly formation of the myelin sheath (10, 11).

The compact myelin sheath is an elaborated multilamellar membrane structure, lipid rich, that envelops selected axons with large diameter. The central nervous system (CNS) myelin is formed by oligodendrocytes windings, while the peripheral nervous system (PNS) myelin is constituted by Schwann cell wrapped around axons. The structure and function of PNS and CNS myelin is similar but differ in composition. In particular, the myelin sheath has several roles including: the control of the propagation speed of action potentials in saltatory conduction, facilitating nerve signal transmission (12–15), and the mechanical support (14, 15). The myelin ultrastructure has a structural unit constituted by the stacking of the four layers: (i) lipidic membrane (lipid polar group, *lpg*), (ii) cytoplasmatic apposition, *cyt*, (iii) a second lipidic membrane, *lpg*, and (iv) extracellular apposition, *ext*, as shown in the pictorial view of Figure 1 (16–18).

**Figure 1:**
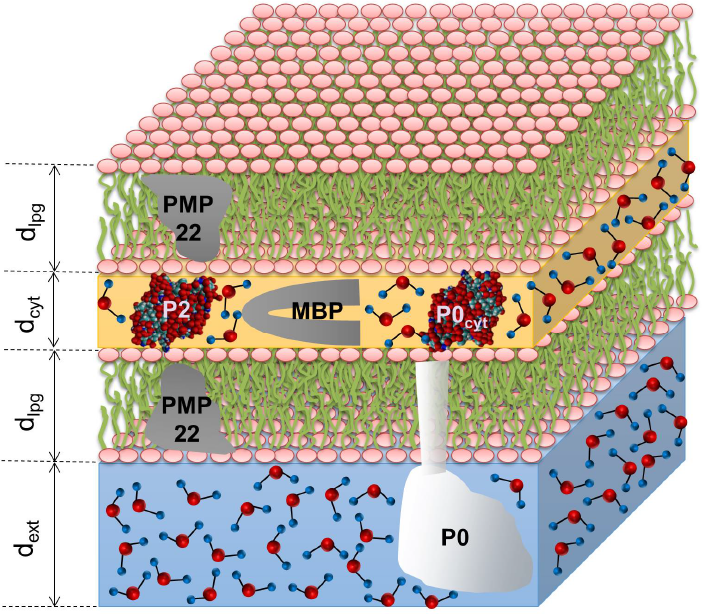
(Color online) Pictorial view of myelin structural unit with the four principal myelin protein of PNS: P0, P2, PMP22 and MBP. The PMP22 protein is located in the membrane, *lpg*. The P0 protein is an integral protein that helps to build the myelin layers stacking. The extracellular domain defines the *ext* layer (blue). The cytoplasmic domain defines the *cyt* layer (yellow), which is the location of the structured protein P2 and the intrinsically disordered protein MBP.

In these systems MBP represent 5-15 % of PNS myelin protein, whereas its abundance is close to 30 % in CNS myelin (14, 15, 19, 20). It is a highly basic, positively charged (15, 21), extrinsic membrane protein present at the cytoplasmic surfaces of compact myelin membranes (*cyt*) (22). MBP is a protein highly soluble in water solvent and it exists in several isoforms with different sequence lengths and charges and there are evidences that it forms dimers (14). The most abundant isoform, in mammalian organisms, has a weight of 18.5 kDa (23). Furthermore, MBP is the principal protein stabilizing the major dense line of CNS myelin, interacting with the negatively charged lipids (10, 14, 24, 25). Through electrostatic interaction with lipid membrane, MBP adopts a more ordered secondary structure (26–28). In PNS myelin it also contributes, together with P0 and P2 proteins (see Figure 1), to compaction of *cyt* (14, 28, 29). Moreover, MBP is a candidate antigen for T-cells and autoantibodies in multiple sclerosis (MS) (15, 30). Recent studies are leading to a new understanding of MBP role as linker and hub in structural and signalling networks in myelin (27, 31–33).

The myelin fluctuations in the out-of-equilibrium functional living state show a correlated disorder that gets lost in the thermodynamic equilibrium obtained by the system degeneration and aging (34–37). Moreover, recently the work of Vattay et al. has proposed that a quantum criticality is at the origin of life predicting biological matter fluctuations (37). The main indicator for proximity to a critical point is the critical opalescence. This is a situation where the size of the different phase regions (ordered, disordered, etc.) begin to fluctuate, in a correlated way, over increasingly large length scales (38), as shown, for instance, in previous work on myelin (17, 18). In this work, we have been interested to clarify the role of myelin proteins dynamics, in particular of MBP, on the myelin ultrastructure functional fluctuations. To tackle this issue, we have used new powerful non-invasive high spatial resolution techniques developed thanks to the latest generation synchrotron sources, advanced focusing optics and fast acquisition methods (17, 18, 39–42). More specifically, we have followed a methodological strategy consisting in two main steps. In the first step we have used Scanning micro X Ray Diffraction (S*μ*XRD) to measure ultrastructural fluctuations of myelin in both out-of-equilibrium physiological and in-equilibrium non-physiological conditions obtained by changing the pH of myelin environment (17, 18). We paid particular attention to the fluctuations of the cytosolic apposition, where the MBP, P2 and P0 proteins act to preserve myelin structure and functionality. In the second step, we have used Small Angle X-ray Scattering (SAXS) to measure the MBP structural fluctuations at nanoscale in a “free state”, namely in an aqueous solution, both at physiological and at non-physiological conditions, obtained by varying the pH of MBP solution. The results should clarify the mechanism of the functional status of the cytoplasmic myelin layer and in particular the role of MBP in this biological system.

## MATERIALS AND METHODS

### Sample preparation

#### SAXS samples preparation

The solute bovine MBP has been purchased from Sigma-Aldrich and used without further purification. We start the preparation by diluting 5 mg of MBP powder in 0.2 ml of pure water, producing a MBP solution at the 25 mg/ml concentration. The samples are made at different pH: for pH < 7 we use a Citrate Buffer with a concentration of 5 *μ*M; for pH > 7 we use the Trizma buffer of 5 *μ*M concentration. For each sample at the various pHs 20 *μ*l of the MBP solution are taken to be added to 80 *μ*l of buffer solution, obtaining a concentration of 5 mg/ml in 100 *μ*l volume. Two solutions are prepared for each pH. In order to evaluate the actual pH of the sample, a laboratory tests are carried out. The samples have been called with the actual pH name obtained.

#### S*μ*XRD samples preparation

The experimental methods were carried out in accordance with the approved guidelines. Adult female frogs (Xenopus Laevis; 12 cm length, 180-200 g weight, Xenopus express, France) were housed and euthanized at the Grenoble Institute of Neurosciences with kind cooperation of Dr. Andre Popov. The local committee of Grenoble Institute of Neurosciences approved the animal experimental protocol. The frogs were individually transferred in water to a separate room for euthanasia that was carried out using a terminal dose of tricaine (MS222) by immersion, terminal anaesthesia was confirmed by the absence of reflexes. Death was ensured by decapitation. Two sciatic nerves were ligated with sterile silk sutures and extracted from both thighs of freshly sacrificed Xenopus frog at approximately the same proximal-distal level through a careful dissection of the thigh. After dissection, the sciatic nerves were equilibrated in culture medium at pH 7.3 for at least 3 hours at room temperature. The culture medium was a normal Ringer’s solution, containing 115 mM NaCl, 2.9 mM KCl, 1.8 mM CaCl_2_, 5 mM HEPES (4-2-hydroxyethyl-1-piperazinyl-ethanesulfonic). Following equilibration, one of the freshly extracted nerves was immediately placed in a thin-walled quartz capillary, 1 mm diameter, sealed with wax and mounted perpendicular to the sample holder, for the SμXRD imaging measurements. In total we acquired 20301 X-ray diffraction patterns, for the fresh sample.

The other frog sciatic nerve, after equilibration, were placed in a solution at pH 6 at room temperature. The culture acidic solution was an acetate buffer solution, made up by mixing 847 ml of 0.1 M acetic acid (CH_3_COOH) and 153 ml of 0.1 M sodium acetate tri-hydrate. Following equilibration, the nerve was placed in a thin-walled quartz capillary, sealed with wax and mounted on the sample holder for a third S*μ*XRD imaging session. In total we acquired 10201 X-ray diffraction patterns, for the denatured sample at pH 6.

A similar procedure was used for the pH 8.5 sample. Another frog sciatic nerve, after equilibration, was placed in a solution at pH 8.5 at room temperature. The culture basic solution was a Tris buffer solution, made up by mixing 100 ml 0.1 M Tris (hydroxymethyl) aminomethane and 29.4 ml of 0.1 M HCl. Following equilibration, the nerve was placed in a thin-walled quartz capillary, sealed with wax and mounted on the sample holder for a third S*μ*XRD imaging session. In total we acquired 10201 X-ray diffraction patterns, for the denatured sample at pH 8.5.

### Experimental and data analysis

#### Small Angle X-ray Scattering

SAXS measurements were performed on the BioSAXS beamline (BM 29) at ESRF (Grenoble, France), equipped with a 2D detector (Pilatus 1M, Dectris). The sample to detector distance for normal operation is 2.867 m, which allows a momentum transfer of *s* = (4*π* sin(*θ*)/*λ*) in the range from 0.05 to 5.8 nm-1. A volume of 50 *μ*l of solution has been placed in a 1.8 mm diameter quartz capillary (mounted in vacuum) with a few tens of micrometer wall thickness, using an automated sample loader developed by EMBL in collaboration with ESRF.

The potential effect of radiation damage has been evaluated performing a 25 s exposure at constant temperature (293 K) without observing any radiation damage. In the experiment, we have used an exposure time of 3 s at each temperature to avoid a possible radiation damage. The sample was initially at room temperature (293 K), and the transfer of the sample to the measurement cell took place at the same temperature.

The data analysis of the MBP SAXS data has been performed exploiting the powerful EOM software package (43), which takes properly into account the coexistence of multiple MBP conformations in solution.

#### Scanning micro X-ray Diffraction

The experimental methods were carried out in accordance with the approved guidelines. The Scanning micro X ray Diffraction measurements of myelin of frog’s sciatic nerve were performed on the ID13 beamline of the European Synchrotron Radiation Facility, ESRF, France. The source of the synchrotron radiation beam is a 18 mm period in vacuum undulator. The beam is first monochromatized by a liquid nitrogen cooled Si-111 double monochromator (DMC) and then is focused by a Kirkpatrick-Baez (KB) mirror system. This optics produces an energy X-ray beam of *λ*=12.6 keV on a 1×1 *μ*m2 spot. The sample holder hosts the capillary-mounted nerve with the horizontal (y) and vertical (z) translation stages with 0.1 *μ*m repeatability. The sample was scanned by using a step size of 5 *μ*m in both y and z direction, in order to avoid a sample damaging and autocorrelation between measured points. A Fast Readout Low Noise (FReLoN) camera (1024×1024 pixels of 100×100 *μ*m2) is placed at a distance of 565.0 mm from the sample to collect the 2-D diffraction pattern in transmission. Diffraction images were calibrated using silver behenate powder (AgC_22_H_43_O_2_), which has a fundamental spacing of d_001_=58.38 Å. We choose an exposure time of 300 ms to minimize radiation damage without affecting sensitivity at the same time (17, 18). The crossed bundle is of approximately 50 myelinated axons. Therefore, the diffraction frames are an average of these axons. Considering the scale of our problem, this is an acceptable average.

We measured different regions of interest (ROIs) in the central part of the nerves around their axis to minimize the capillary geometry effect on the X-ray absorption. A typical 2-D diffraction pattern of myelin shows the expected arc-rings corresponding to the Bragg diffraction orders h = 2, 3, 4, 5. The 2D diffraction patterns have been radially integrated to provide 1D intensity profiles, *I* (*s*), vs. transfer moment *s* = 2*π* sin(*θ*)/*λ*, after background subtraction and normalization with respect to the impinging beam. The 1D radial profile shows the four characteristic peaks of myelin modelled with a Lorentian line shape from which we get the peak amplitude, *A*(*h*), and full width at half height, *w*(*h*), used for the Fourier’s analysis yielding electron density for each pixel according to:

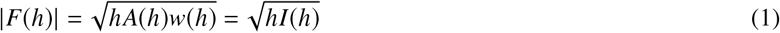

for each reflection of order *h*. These structure factors, |*F*(*h*)|, were employed in a Fourier analysis to extract the Electron Density Distribution (EDD) of myelin:

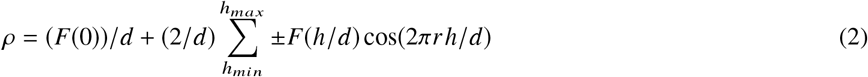

where the phases were taken from literature (16, 44):

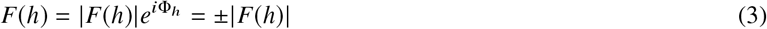

since the nerve is a centrosymmetric structure, so we consider just the real terms of Fourier series.

From the differences between two adjacent maxima in the EDD profile the widths of the inter-membrane spaces at the cytoplasmic (d_*cyt*_) and extracellular (d_*ext*_) appositions and the thickness of the lipid bilayer (d_*lpg*_) were obtained. From these we can obtain the period of the structural unit d_*λ*_=2d_*lpg*_+d_*ext*_+d_*cyt*_ (16, 44). The extrapolation of EDD at each pixel of the ROI, was performed using a customized in-house developed code written in MatLab (Mathworks, Natick, MA).

## RESULTS AND DISCUSSION

### Myelin ultrastructural dynamics as a function of pH

As already said above, the myelin ultrastructure is constituted by a repetition of a structural unit, made by the stacking of four sub-layers: *cyt*, *lpg*, *ext* and another *lpg* (16–18) (see Figure 1). This structure gives rise to well-known XRD Bragg peaks. The period and sublayer thicknesses, labelled d_*λ*_, d_*cyt*_, d_*lpg*_, d_*ext*_, have been extracted from the electron density distribution of each sublayer, computed by Fourier analysis of the XRD profiles, as described in detail elsewhere (17, 18) and in the Methods section. Thus, we have built the spatial maps for each thickness and for each map we have calculated the probability density functions (PDF) for acid pH 6 and basic pH 8.5 samples, both values around the functional pH 7.3 state. This range of pH values is physiological, i.e. values of pH within this range are those allowed by a living system in trauma conditions, such as an ictus; larger variation of pH are not physiologically tolerable. Another important parameter, besides PDF, to describe quantitatively the myelin ultrastructural dynamics, is given by the ratio *ξ* = (*d*_*ext*_ + *d*_*cyt*_)/d_*lpg*_ introduced by Campi et al. (18). Indeed, this parameter allows us to characterize the overall myelin state uniquely. The following considerations are supported by several statistical quantities such as mean value, standard deviation and relative standard deviation (RSD) used to quantify myelin ultrastructure fluctuations (see Table 1).

**Table 1:** Statistical analysis of the of the thickness d_*cyt*_, d_*ext*_, d_*lpg*_ and d_*λ*_, in the fresh and denatured with pH=6 and pH=8.5 samples, from the electron density profiles for Xenopus Laevis sciatic nerves. Here we report mean values, *μ*, standard deviations, *σ*, and Relative Standard Deviation RSD = *σ*/*μ*.

| samples | $\mu(\text{nm})$ | | | $\sigma(\text{nm})$ | | | RDS(%) | | |
| --- | --- | --- | --- | --- | --- | --- | --- | --- | --- |
|  | fresh | pH=6 | pH=8.5 | fresh | pH=6 | pH=8.5 | fresh | pH=6 | pH=8.5 |
| $d_{cyt}$ | 2.999 | 3.105 | 3.123 | 0.036 | 0.094 | 0.053 | 1.20 | 3.04 | 1.69 |
| $d_{ext}$ | 4.860 | 4.719 | 4.711 | 0.082 | 0.362 | 0.141 | 1.68 | 7.67 | 2.99 |
| $d_{lpg}$ | 4.746 | 4.755 | 4.800 | 0.055 | 0.224 | 0.081 | 1.15 | 4.72 | 1.70 |
| $d_\lambda$ | 17.350 | 17.362 | 17.468 | 0.024 | 0.059 | 0.047 | 0.14 | 0.34 | 0.27 |

In Figure 2a we show the PDF of the d_*λ*_, d_*cyt*_, d_*lpg*_, d_*ext*_ thicknesses for the acid pH 6 sample. The dashed lines represent the PDFs of the same quantities calculated for the functional sample (pH 7.3), as described in Campi et al. (18), to be used as reference. Despite the broadened distribution, the period, d_*λ*_, maintains nearly constant its average value. In particular, we find that the RSD for d _*λ*_ is 0.14% in the fresh sample while reaches a value of 0.34% for the pH=6 sample. On the other hand, we can note that the average values of d_*cyt*_ and d_*lpg*_ increase, while d_*ext*_ decreases. This spatial negative correlation between hydrophobic lipidic membrane and hydrophilic extracellular apposition ensures the stability of the period, as showed in Table 2 where the Pearson’s correlation coefficients are reported. This negative correlation can be clearly visualized in Figure 2b by the scatter plot of *ξ* as a function of the period d_*λ*_, the dispersions 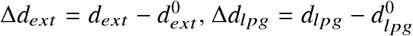 and 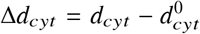, where 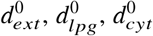 are the reference values for protein-free myelin, given by 2.2, 3.7, 2.2 nm (28), respectively. The grey solid circles representing the fluctuations found in the fresh nerve are reported for comparison. In this acidic case the PDF of denatured sample, shown in the left panel of Figure 2b, broadens dramatically in comparison with the situation at physiological state at pH 7.3. Indeed, we notice the non-zero probability to have the conformational parameter larger than 1, sign of degeneration of the sheath, due to greater proton density (45–47) and myelin protein denaturation. The PDF of denatured sample cannot be fitted because it does not show a clear analytical shape and loses the fat tail of the Levy distribution describing the correlated disorder of the functional out-of-equilibrium state (18, 48, 49).

**Figure 2:**
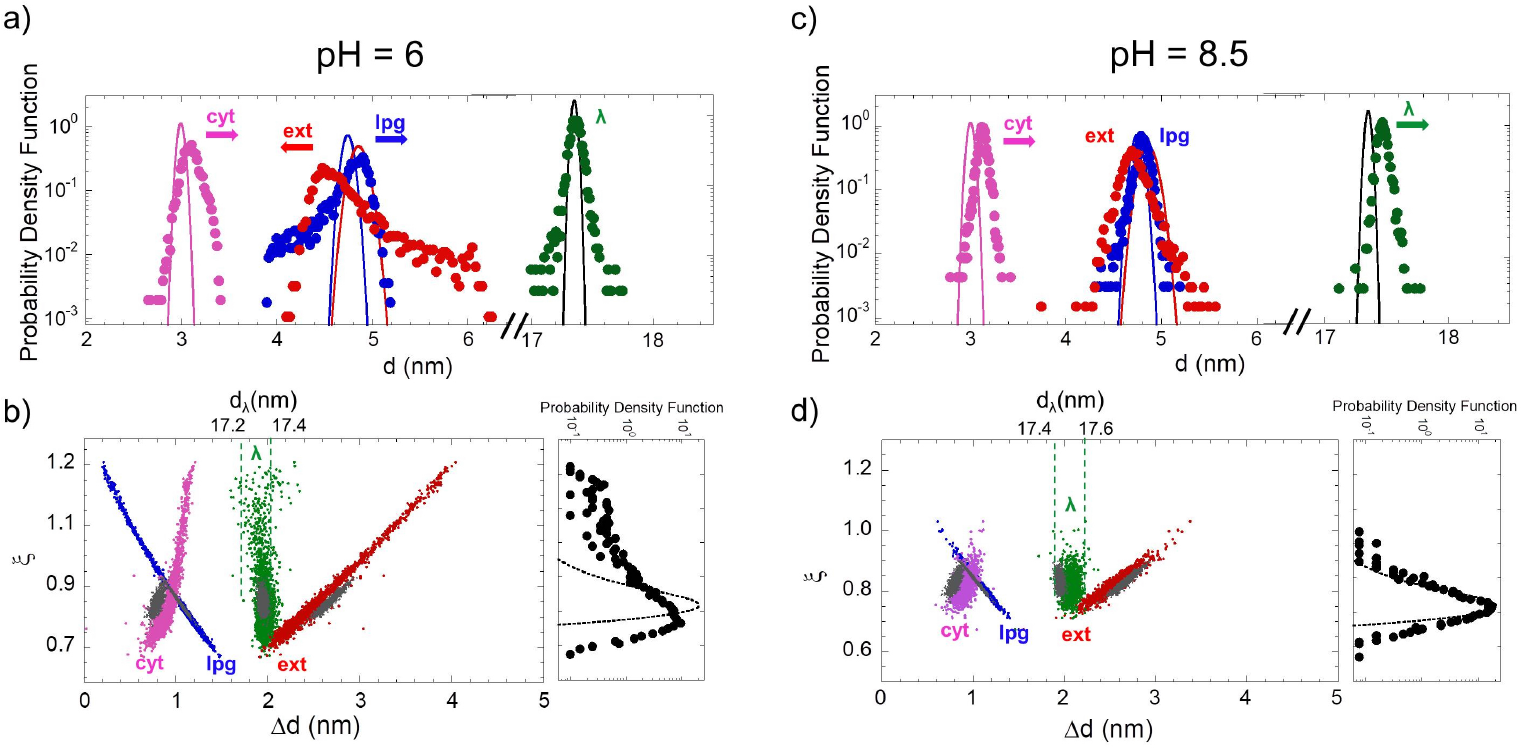
(Color online) a) The probability density functions (PDFs) of the thicknesses d_*λ*_, d_*cyt*_, d_*lpg*_, d_*ext*_ of the different myelin layers for acid pH=6 sample. The continuous lines represent the PDFs of the fresh functional sample at pH=7.3, for comparison. b) Scatter plot of the myelin conformational parameter, *ξ*, as a function of the deviations Δ*d*_*ext*_, Δ*d*_*lpg*_ and Δ*d*_*cyt*_ of the layer thicknesses from the protein-free membrane (28). The plot shows the anticorrelation between the lipidic membrane and extracellular apposition. *ξ* as a function of the period, d_*λ*_, (top x-axis) is reported between two dotted lines at d_*λ*_=17.2 nm and d_*λ*_=17.4 nm. In the right panel, the PDF of *ξ* illustrates the crossover from the functional state (dashed line) to the degraded state at acidic pH (full circles). The denatured PDFs isn’t fitted because shows very large incoherent fluctuations at high values of the conformational parameter *ξ*. c) PDFs of d_*λ*_, d_*cyt*_, d_*lpg*_, d_*ext*_ for basic pH 8.5 sample. Here the continuous lines represent always the PDFs of the fresh functional sample, for comparison. d) Scatter plot of the conformational parameter, *ξ*, as a function of Δ*d*_*ext*_, Δ*d*_*lpg*_ and Δ*d*_*cyt*_ and d_*λ*_. The plot shows the anticorrelation between the lipidic membrane and extracellular apposition as in the acidic case. In the right panel we show the Probability density function of *ξ* in the pH 8.5 sample which show minor fluctuations of conformational parameter, in comparison with the acidic case, as reported in Table 1.

In Figure 2c we show the PDF for the d_*λ*_, d_*cyt*_, d_*lpg*_, d_*ext*_ thicknesses, resulting from the same analysis described above, but this time applied to the myelin sample at basic pH 8.5. Also in this plot the dashed lines represent the PDFs of the same quantities calculated in the functional sample, as reference. At this pH value we can notice a slight increase of period compared with the physiological sample, due to the fact that the increase of d_*lpg*_ and d_*cyt*_ is no more compensated by a corresponding decrease of d_*ext*_. Although the observed increase of the fluctuations of the sublayers thicknesses with respect to the functional sample (see Table 1), these fluctuations are quite smaller in comparison with those observed for the acid pH 6 sample. It is due to the larger proton density in the acidic sample that gives a worse resistance of myelin PSN protein structure in comparison with the sample at basic pH (15, 21). The smaller fluctuations in the basic pH 8.5 sample are well depicted both i) in the scatter plot of Figure 2d, showing *ξ* as a function of the period d_*λ*_, Δ*d*_*ext*_, Δ*d*_*lpg*_ and Δ*d*_*cyt*_ and ii) in the denatured sample PDFs of *ξ* where we observe just a slight deviation from the physiological case. Thus, for our aims, in order to investigate the relationship between the myelin fluctuations and protein dynamics in the transition from out-of equilibrium to in-equilibrium conditions, we’ll focus on the degeneration of the system induced by setting the pH of the system at acidic conditions.

**Table 2:**
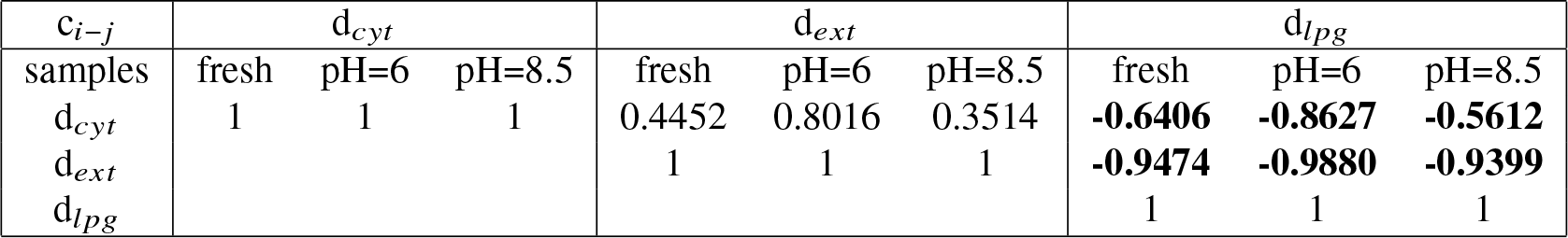
Correlation coefficients c_*i−j*_ between maps of the spatial fluctuations between the different layers i, j = *cyt*, *ext* and *lpg* in the fresh, pH=6 and pH=8.5 samples. Negatively correlated coefficients are indicated by bold. These correlations are well explained by Figure 2 for each sample.

| $c_{i-j}$ | $d_{cyt}$ | | | $d_{ext}$ | | | $d_{lpg}$ | | |
| --- | --- | --- | --- | --- | --- | --- | --- | --- | --- |
| samples | fresh | pH=6 | pH=8.5 | fresh | pH=6 | pH=8.5 | fresh | pH=6 | pH=8.5 |
| $d_{cyt}$ | 1 | 1 | 1 | 0.4452 | 0.8016 | 0.3514 | **-0.6406** | **-0.8627** | **-0.5612** |
| $d_{ext}$ | | | | 1 | 1 | 1 | **-0.9474** | **-0.9880** | **-0.9399** |
| $d_{lpg}$ | | | | | | | 1 | 1 | 1 |

### “Free” MBP structure by SAXS measurements

In the second step of our study, we have investigated the MBP structural fluctuations at room temperature in the “free state”, that is an MBP aqueous solution at the physiological value of pH 7.3 and at two pH values around it, namely at pH 6 and pH 8.5. Moreover, MBP solution at pH 5 and pH 4 were also investigated, to get a complete structural information far from the living state (18). We have used synchrotron X-ray Small Angle Scattering (SAXS) allowing to inspect MBP size, dimension and protein shape at nanoscale.

MBP SAXS profiles, I(q), in the four different acid pH with the same buffer solution and their fit (green line) obtained by means of the Ensemble Optimization Method (EOM) are shown in Figure 3a. EOM first generates a pool of models spanning the protein’s conformational space (50000 conformations in our case), based upon sequence and structural information; then selects from this pool an ensemble of conformers that best fits the SAXS experimental data, by using an iterative genetic algorithm (3). Upon inspection of the I(q) curves at the different pH values investigated, no significant differences between these curves can be observed. Indeed, the radius of gyration, R_*g*_, extracted by Guiner approximation and shown in Figure 3b, does not change significantly with pH, following a constant trend, typical of intrinsically disorder protein behavior (50). In Figure 3b there are also reported the values of cytosolic apposition for PNS at physiologic condition, PNS after denaturation at pH 5 and CNS at physiological condition (16, 18, 51). These values show that the MBP in myelin is always in a very compressed state, compared to the free state, considering also that in the *cyt* myelin layer the diameter of MBP should be considered and not its radius. Furthermore, we can notice that as the pH decreases the standard deviation around the R_*g*_ mean values decreases, indicating a larger rigidity of structure at low pH, or, in other words, an increased ability of protein to assume more conformations in physiological conditions (52). In Figure 3c we represent the 3D structure of the six EOM conformations of MBP, used to obtain the SAXS fit in Figure 3a. Each conformation has specific values of R_*g*_, maximum size, D_*max*_, and percentage weight used in the fitting algorithm. The MBP probability density function (PDF) of R_*g*_ and D_*max*_ calculated from these six EOM conformers are shown with red symbols in Figure 3d and 3e, respectively. Here we also show the PDFs obtained by the Fourier transform of the SAXS patterns, with black symbols. The colored arrows correspond to the R_*g*_ and D_*max*_ values of a single conformation shown in panel c of Figure 3, using the same color code. Our EOM analysis suggests that MBP can perform large functional fluctuations assuming conformations whose size, i.e. R_*g*_, approaches small values down to around 2 nm and large values up to 7 nm. All these information obtained are useful to comprehend MBP role in myelin functional fluctuations and compaction.

**Figure 3:**
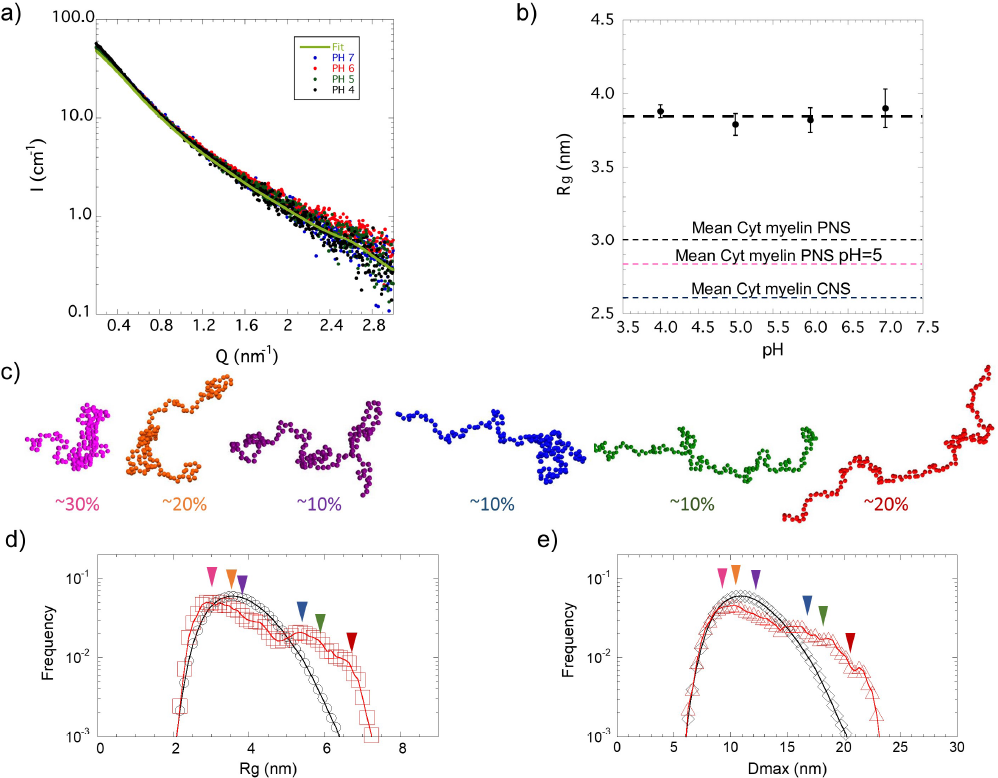
(Color online) a) SAXS profiles measured at room temperature of the intrinsically disordered protein MBP in aqueous solution at different pH alongside the curve fit (green line) obtained by Ensemble Optimization Method (EOM) analysis. b) The MBP radius of gyration (R_*g*_) as a function of the pH. The dashed lines indicate the mean values of cytosolic apposition of PNS at physiological condition, PNS after denaturation at pH 5 and CNS at physiological condition. c) A 3D representation of six possible conformations of MBP extracted by EOM, used to fit the SAXS data. Their percentages used in the fit algorithm are reported. Probability density function of d) R_*g*_ and of e) maximum size, D_*max*_, of MBP in solution obtained by EOM (red symbols) using the six EOM conformations and standard Fourier Transform (black line) of SAXS profiles. The arrows with different colors indicate each contribution at R_*g*_ and D_*max*_ corresponding to the colors of the 3D protein representations of panel c).

### MBP and structured proteins dynamics in cytosolic layer of myelin

We move now to investigate the degeneration of the cytoplasmic apposition, obtained by sample denaturation at acid pH 6 and pH 5 (18). The PDFs of d_*cyt*_ for the pH 6 (pink circles) and pH 5 (magenta circles) samples, compared with the functional sample at pH 7.3 (dashed line) are shown in Figure 4a, along with the mean values of d_*cyt*_ for the physiological and pH 5 samples indicated by a magenta and black vertical line, respectively. The allowed values of the CNS cytoplasmatic apposition ranges from 2.50 to 2.95 nm as indicated by the band of violet shades (51, 53).

**Figure 4:**
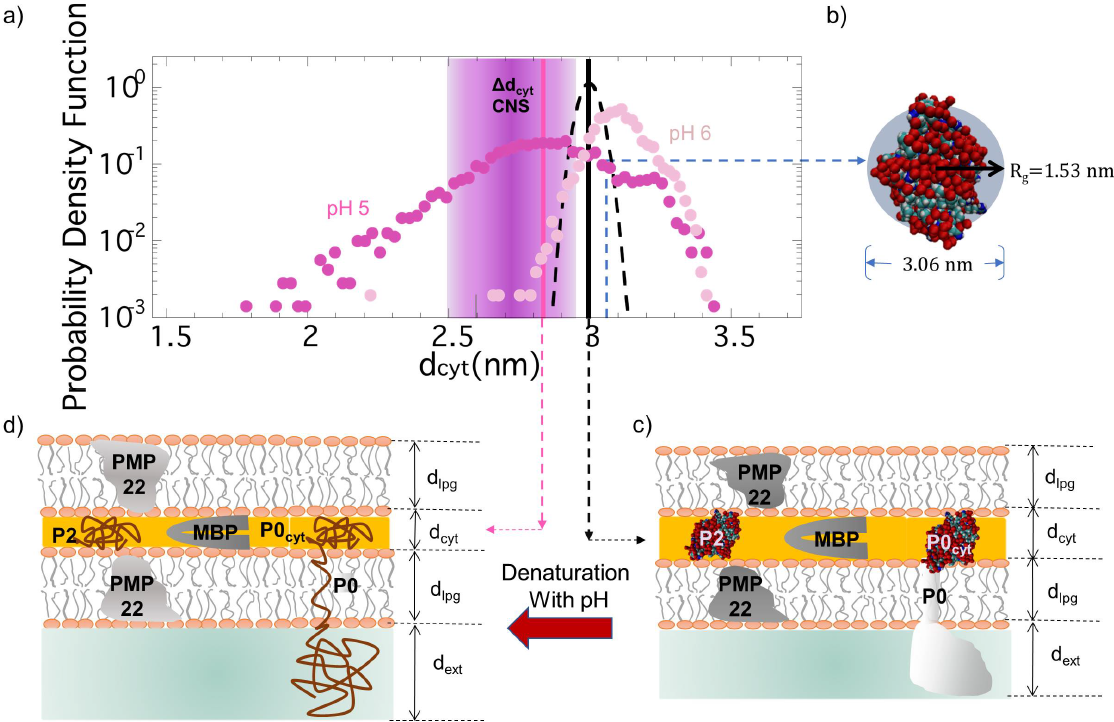
(Color online) a) Probability Density Function of the cytosolic apposition thickness, *d*_*cyt*_, at pH 5, in comparison with samples at pH 7.3 and pH 6 in semi-log plot. The magenta full circles represent the PDF of *d*_*cyt*_ at pH 5, compared with the PDF of *d*_*cyt*_ in the samples at pH 6 (pink full circles) and pH 7.3 (dashed line). The solid vertical lines indicate the mean values of the physiological (black) and pH 5 (magenta) samples. The violet shaded area indicates the allowed values for the cytosolic apposition in the central nervous system (CNS). b) A 3D representation of P2 folded structure. Pictorial view of myelin structural unit in fresh (c) and denatured sample at pH=5 (d), where the structured proteins P2 and P0 became unfolded with compression of the cytosolic thickness while the MBP continues to display large functional biological fluctuations.

We can see that the PDF widens as the pH decreases. In particular, cytoplasmic fluctuations increase as the pH value moves away from the physiological value, while their mean values have a different behavior. At pH 6 the d_*cyt*_ mean value is slightly larger than that of functional sample, whereas for the pH 5 sample is slightly shorter. Correspondingly, the fluctuations RSD increase dramatically, going from 1.2 % in the fresh sample to 3.04 % and 8.37 % for the pH=6 and pH=5 samples, respectively. For the pH 6 sample, the little increase of d_*cyt*_ spacing are consistent with the maintained structure of the myelin proteins P2 and P0, as their R_*g*_ (1.53 nm and 2.2 nm) (54, 55), together with MBP action, which has increased its rigidity, therefore its strength (Figure 3b). On the other hand, for the pH 5 sample the P2 and P0 structures become unfolded because the *cyt* apposition fluctuates assuming values too small, no longer consistent with the size of structured and folded proteins. Therefore, the *cyt* layer thickness is maintained by the only functioning protein, MBP, and its mean value falls within the possible values for that of the CNS (Figure 4a) (51, 53), made up only by MBP (14). In particular the cytoplasmic thickness reaches values down to 2 nm. It is interesting to notice that this latter value corresponds to the limit value for anomalous diffusion of water between two hydrophilic surfaces (48, 56–61) and also to the smallest value of the PNS myelin cytoplasmic apposition (28). A 3D representation of P2 folded structure is shown in Figure 4b. Thus, in the functional out-of-equilibrium state, the structured proteins P0 and P2 act as pins, maintaining the cytosolic apposition anchored to around its physiological value (d_*cyt*_ = 3 nm). Around this value, small cytosolic physiological fluctuations occur, thanks to the MBP protein that fluctuates assuming compact conformations, as found in our SAXS measurements (see Figure 3). The pictorial view of myelin ultrastructural unit in Figure 4c schematizes the situation in the out-of-equilibrium physiological state. Moving towards in-equlibrium conditions, at low pH 5, the large fluctuations measured for the cytosolic apposition are due to the denaturation of the structured proteins that remove the structural hinges for the cytosol. However, despite these large fluctuations, the *cyt* layer preserves its structure, and continues to stay in a still functioning condition, due to the action and function of the MBP, whose size fluctuates stably from 2 to 7 nm, given its intrinsically disordered nature. Figure 4d represents the situation resulting from denaturation of pH 5, showing the cytosolic compression. This scenario suggests that the most significant alterations with respect to the functional state of PNS myelin depend mainly from the structural degeneration of the *lpg* membrane and of the *ext* layer.

## CONCLUSION

The correlated disorder found in the functional fluctuations o f myelin ultrastructure constitutes a recent point of high interdisciplinary interest involving fundamental statistical and biological physics. In this context the study of the myelin protein dynamics provides relevant insight in the understanding the biological functionality with a major completeness in a simple model system. Here together with structured proteins, P0, P2 working in the cytosol, and the PMP22 in the lipid membrane, we have the intrinsically disordered protein MBP acting in the cytosol. Despite the classical structure-function paradigm, where a unique biological function is defined by its specific structured state determined by the amino acid sequence, the IDP are assuming more and more relevance thanks to the fact that their conformational flexibility and structural plasticity play a vital role in many biological processes. In this work we have investigated the interplay of myelin ultrastructural fluctuations and MBP dynamics in the transition from out-of-equilibrium functional state to the in-equilibrium degraded state. At this aim we have used Scanning micro X Ray Diffraction and Small Angle X Ray Scattering to measure the fluctuations of myelin ultrastructure and MBP, respectively. We did it both in physiological out-of-equilibrium and degraded in-equilibrium conditions obtained by changing the pH of the system. More specifically, we focused on the acidic pH since here we got larger deviations from the physiological state in comparison with the basic pH. We have found that in the physiological state the cytosolic layer shows small fluctuations around the value of 3 nm, given by the size of the P2 and P0_*cyt*_ proteins, acting as pins. The entity of these small fluctuations of d_*cyt*_ is ensured by the MBP that assumes compact conformations, around 3 nm in synergy with structured P2 and P0 protein dynamics. Moving towards in-equilibrium, degraded conditions, at low pH, the cytosol fluctuates from 2 to 7 nm. Approaching these extreme values, the structured proteins P2 and P0 unfold and lose their pin function, while the flexibility of the intrinsically disordered MBP allows to follow these large fluctuations, preserving the cytosol structure and functionality. Therefore, the most significant alterations from the functional state of PNS myelin depend mainly by the structural degeneration of the *lpg* and of the *ext* layers. In conclusion, the present study highlights further and unknown aspects of degeneration of PNS myelin, adding information about the comprehension of this problem both in neurodegenerative diseases and those resulting from a trauma.

## AUTHOR CONTRIBUTIONS

A. Bianconi and G. Campi designed the S*μ*XRD experiment. M. Burghammer provided the XRD station at ESRF; M. Di Gioacchino, G. Ciasca and G. Campi designed the SAXS experiment; G. Campi, M. Di Gioacchino, G. Ciasca and A. Bianconi performed the data analysis; and M. Di Gioacchino, G. Campi, F. Bruni and A. Bianconi wrote the paper.

## ACKNOWLEDGMENTS

We are grateful to the staff of ID13 and BM29 beamlines at ESRF for experimental help. The authors thanks Dr. Andre Popov at the G.I.N. for animal housing and specimen preparation. We thank T. A. Hawkins of the Department of Cell and Developmental Biology at UCL London for experimental help in the early stage of the experiment. We thank superstripes-onlus for financial support. G.C. acknowledges the support of the Institute of Crystallography of Consiglio Nazionale delle Ricerche. The authors acknowledge useful discussions with N. Poccia and A. Ricci.

